# Evolutionary model of protein secondary structure capable of revealing new biological relationships

**DOI:** 10.1101/563452

**Authors:** Jhih-Siang Lai, Burkhard Rost, Bostjan Kobe, Mikael Bodén

**Author notes:** Corresponding author *Email addresses* (Jhih-Siang Lai), (Mikael Bodén).

## Abstract

Ancestral sequence reconstruction has had recent success in decoding the origins and the determinants of complex protein functions. However, phylogenetic analyses of remote homologues must handle extreme amino-acid sequence diversity resulting from extended periods of evolutionary change. We exploited the wealth of protein structures to develop an evolutionary model based on protein secondary structure. The approach follows the differences between discrete secondary structure states observed in modern proteins and those hypothesised in their immediate ancestors. We implemented maximum likelihood-based phylogenetic inference to reconstruct ancestral secondary structure. The predictive accuracy from the use of the evolutionary model surpasses that of comparative modelling and sequence-based prediction; the reconstruction extracts information not available from modern structures or the ancestral sequences alone. Based on a phylogenetic analysis of multiple protein families, we showed that the model can highlight relationships that are evolutionarily rooted in structure and not evident in amino acid-based analysis.

## 1. Introduction

Zuckerkandl and Pauling’s idea of inferring ancestors from modern pro­teins (Zuckerkandl and Pauling, 1965) has had recent success in decoding the origins and the determinants of complex protein functions (Hochberg and Thornton, 2017). For example, by comparing inferred ancestors to modern proteins, Wilson et al. (2015) elucidated binding affinities in protein kinases, Clifton and Jackson (2016) decoded origins of promiscuity in a solute-binding protein family, and Hudson et al. (2016) identified historical amino acid substitutions that alter specificity of a DNA-binding domain. Ancestral sequence reconstruction typically requires a phylogenetic tree, multiple modern protein sequences in an alignment, and an evolutionary model that assigns a probability to evolutionary events. Ancestral sequence reconstruction establishes plausible historical protein sequences by searching the most probable mutation events to bring about the composition displayed by modern proteins.

The point accepted mutation (PAM) matrix was the first successful model to describe evolutionary changes for amino acids (Dayhoff et al., 1978). Dayhoff, Schwartz and Orcutt defined the probabilities of substitution between amino acid states and each state’s stationary probability, based on counts of implied change among homologous proteins with high sequence identities. Jones, Taylor and Thornton broadened Dayhoff’s approach to work with larger data-sets (Jones et al., 1992). Recent approaches to generate evolutionary models often extend empirical models by using them as starting points and adjust them to optimally recover modern protein sequences irrespective of their similarity (Whelan and Goldman, 2001). A number of improvements have increased the robustness of Dayhoff’s original idea (Kosiol and Goldman, 2005).

While only indirectly subject to mutation, a protein molecule alters its three-dimensional shape to explore and select biological function over evolutionary time. A significant body of research has established that structural features are strongly linked to evolution (Worth et al., 2009; Echave et al., 2016; Kinch and Grishin, 2002; Pal et al., 2006). Goldman et al. (1998) explored this close relationship; they estimated amino acid models specific to each class of secondary structure, and incorporated the collective of models into a Markov chain model of amino acid replacement. This model is capable of indirectly considering secondary structure in phylogenetic analyses. Le and Gascuel (2010) similarly created specific amino acid models for secondary structure classes and solvent accessibility.

The present work takes advantage of the wealth of protein structures in the Protein Data Bank (PDB) (Rose et al., 2017) to develop an evolutionary model that *explicitly* treats secondary structure states as evolutionary traits; we explore a novel basis to detect homologous relationships. We establish that phylogenetic analyses of protein families with this model are biologically meaningful in their own right; they also add value to analyses based on an amino acid-based evolutionary model. The model promises to enable phylogenetic analyses of large families with extreme sequence diversity. To demonstrate the ability to guide phylogenetic analysis and the utility as a biologically meaningful metric, we developed methods to infer phylogenetic trees, and to reconstruct ancestral secondary structure.

## 2. Results

### 2.1. Evolutionary model of secondary structure

We used Dayhoff’s original approach (Dayhoff et al., 1978) and that of Jones, Taylor and Thornton (1992) to explore evolutionary models of secondary structure states; collectively, we refer to this class of models as Evolu-sec. We refer to approach-specific models by author and data type, e.g., DSO-sec and JTT-sec are secondary structure models proposed in this work based on Dayhoff, Schwartz and Orcutt’s approach and Jones, Taylor and Thornton’s approach, respectively; DSO (Dayhoff et al., 1978) and JTT (Jones et al., 1992) are published amino acid evolutionary models. We framed the resulting models as instantaneous rate matrices (IRMs). An IRM contains the rates of substitution of characters indexed *i* (row) by *j* (column) when *i* ≠ *j*; diagonal entries *i* = *j* are defined by the requirement that each row sums to 0 (Liò and Goldman, 1998). IRMs have appealing theoretical properties and are widely adopted, simplifying adaptations of this work to existing phylogenetic tools (Kosiol and Goldman, 2005).

To parameterise our IRMs, we retrieved all protein crystal structures with a resolution <4.0 Å from the PDB. We use DSSP (Kabsch and Sander, 1983) to assign one of seven secondary structure states to each residue, i.e., B (beta-bridge), E (beta-strand), G (helix-3), H (alpha-helix), I (helix-5), S (bend), and T (turn), excepting only those residues that fall outside DSSP’s definitions. How data are used depends on approach and results in slightly different IRMs; our default IRM is based on the DSO-approach and is shown in Table 1. The precise steps of the approaches and the properties of the models are described in the Supporting Information (SI) appendix, and data sets are provided in Additional Data Tables S1-S2.

**Table 1:** Instantaneous rate matrix for secondary structure based on Dayhoff’s approach (DSO-sec; values are multiplied by 10,000).

|  | B | E | G | H | I | S | T |
| --- | --- | --- | --- | --- | --- | --- | --- |
| B (Beta-bridge) | -23678 | 17919 | 0 | 0 | 0 | 4801 | 958 |
| E (Beta-strand) | 634 | -1946 | 66 | 19 | 0 | 995 | 232 |
| G (Helix-3) | 0 | 402 | -28426 | 7819 | 0 | 3317 | 16889 |
| H (Alpha-helix) | 0 | 14 | 993 | -5119 | 60 | 528 | 3523 |
| I (Helix-5) | 0 | 0 | 0 | 263624 | -441405 | 0 | 177782 |
| S (Bend) | 530 | 3104 | 1703 | 2136 | 0 | -20112 | 12638 |
| T (Turn) | 73 | 500 | 5986 | 9831 | 113 | 8721 | -25224 |

### 2.2. Secondary structure-based phylogenetic inference

We extracted 1,591 domain entries from the Pfam database (Finn et al., 2016), providing us with curated sequence alignments and corresponding secondary structure assignments based on DSSP. We determined evolutionary distances by maximum likelihood (Felsenstein, 1996) for both secondary structure and amino acid representations of all pairwise alignments, i.e. the distance *t* via a hypothetical ancestor that best explains each pair of sequences.

Based on 22,157 alignments from Pfam, the Pearson correlation between the secondary structure and amino acid distances is 0.54 (Fig. 1 [left]). *t* is scaled to the unit of “expected number of substitutions per site”; the trends in Fig. 1 suggest that protein domains are generally more conserved in secondary structure, relative to amino acid sequence. To understand the potential impact of mis-aligned amino acid sequence pairs, we also determined evolutionary distances from structure superposition of the same pairs using jCE (Prlic et al., 2010) (Fig. 1 [right]).

**Figure 1:**
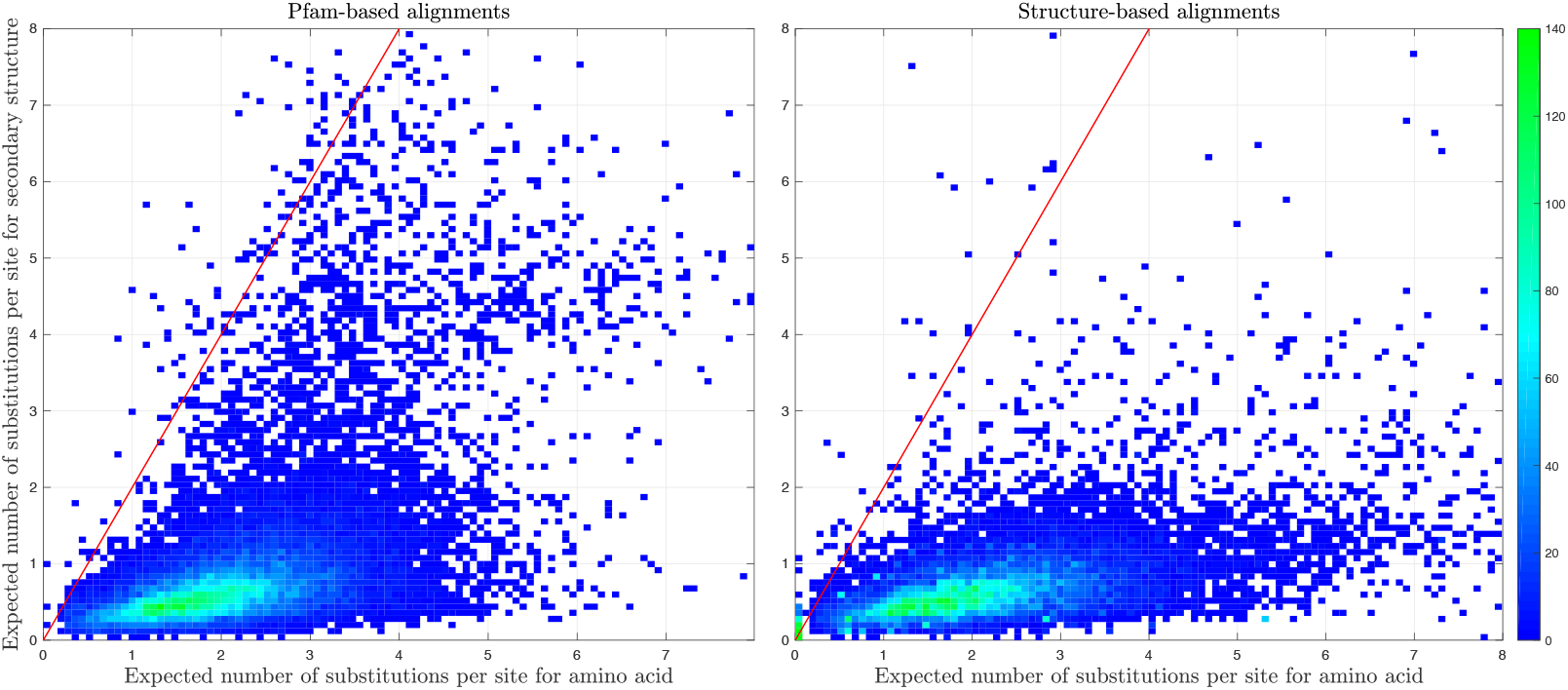
Left: Evolutionary distances by secondary structure and amino acid models (DSO-sec and JTT, respectively) for the same 22,157 alignments as defined in Pfam (left) and by structure superposition (right; using jCE (Prlic et al., 2010)). Each tile in the heatmap represents the number of alignments falling into the same combination of time intervals, i.e. the same numbers of substitutions of amino acids (*x* axis) and secondary structure states (*y* axis).

Based on the original Pfam alignments, 110 pairwise alignments (involv­ing 48 domain entries) have double the expected number of secondary structure relative amino acid substitutions (red line in Fig. 1 [left]). Two domains dominate this set of spuriously aligned homologs: WD40 repeat (PF00400) and haloacid dehalogenase-like hydrolase (PF00702); they form major clusters in the Pfam data set with 14 alignments each. The re-aligned pairs differ substantively from those of Pfam (Fig. 1 [right]): 76 of the 110 pairs have fewer expected substitutions in secondary structure relative to amino acids (see Additional Data Table S3).

Given that the evolutionary models were derived using the same principles, the low correlation is noteworthy. At the very least, this suggests that the two data types on which the models are based are complementary; we propose that inferences done with one will not necessarily capture all aspects of the other. To test if phylogenetic analysis of protein families encoded by secondary structure states leads to biological insights, we implemented a maximum likelihood method to infer a phylogenetic tree based on a multiple sequence alignment given any evolutionary model, including the seven-state model of secondary structure.

We used DSO-sec (as well as JTT-sec and JTT for comparison) to explore evolutionary relationships among members of the Toll/interleukin-1 receptor (TIR) domain family. These domains are found in proteins involved in innate immunity pathways (and related processes) and are present across plants, mammals, bacteria and archaea. TIR domains have similar structures but sequence identities are often less than 20% (Ve et al., 2015). Despite the lack of similarity at the amino acid-level, they share a flavodoxin-like fold with many functionally diverse proteins. In mammals, TIR domains are found in Toll-like receptors (TLRs) and interleukin-1 receptors, as well as adaptor proteins involved in signalling downstream to transcription factors that control the immune response and inflammation (Akira and Takeda, 2004; O’Neill and Bowie, 2007; Boraschi and Tagliabue, 2013). TIR domains in plants are found in nucleotide-binding domain/leucine-rich repeat resistance proteins (NLRs) (Bentham et al., 2017). TIR domains are also found in bacteria, where at least some of them interfere with the host immune system (Rana et al., 2013).

TIR domains function as a homotypic interaction motif, and the molecular basis of interactions has recently been characterized for TLR4 signalling assembly with MyD88 and MAL (Ve et al., 2017). The TIR domains from the mammalian protein SARM1 and some bacterial proteins have also been shown to have NAD^+^-cleavage activity (Essuman et al., 2017, 2018); this finding suggests there may be evolutionary, rather than merely structural links with bacterial enzymes such as the CMP hydrolase MilB. We therefore considered that an evolutionary analysis could provide insights into the functions of TIR domains and structurally similar proteins.

We used data from 64 structures gathered through a Dali search (Holm and Laakso, 2016) to construct three different phylogenetic trees (see SI Figures 2-4, and Additional Data Tables S4-S5). Two trees were based on structure superposition using jCE (Prlic et al., 2010): DSO-sec+jCE (Fig. 2 left) and JTT+jCE (Fig. 2 middle), and one on a classical, amino acid-based alignment: JTT+MAFFT (Fig. 2 right). Since the JTT-sec+jCE tree has similar topology to DSO-sec+jCE we compared the trees generated by DSO-sec and JTT.

**Figure 2:**
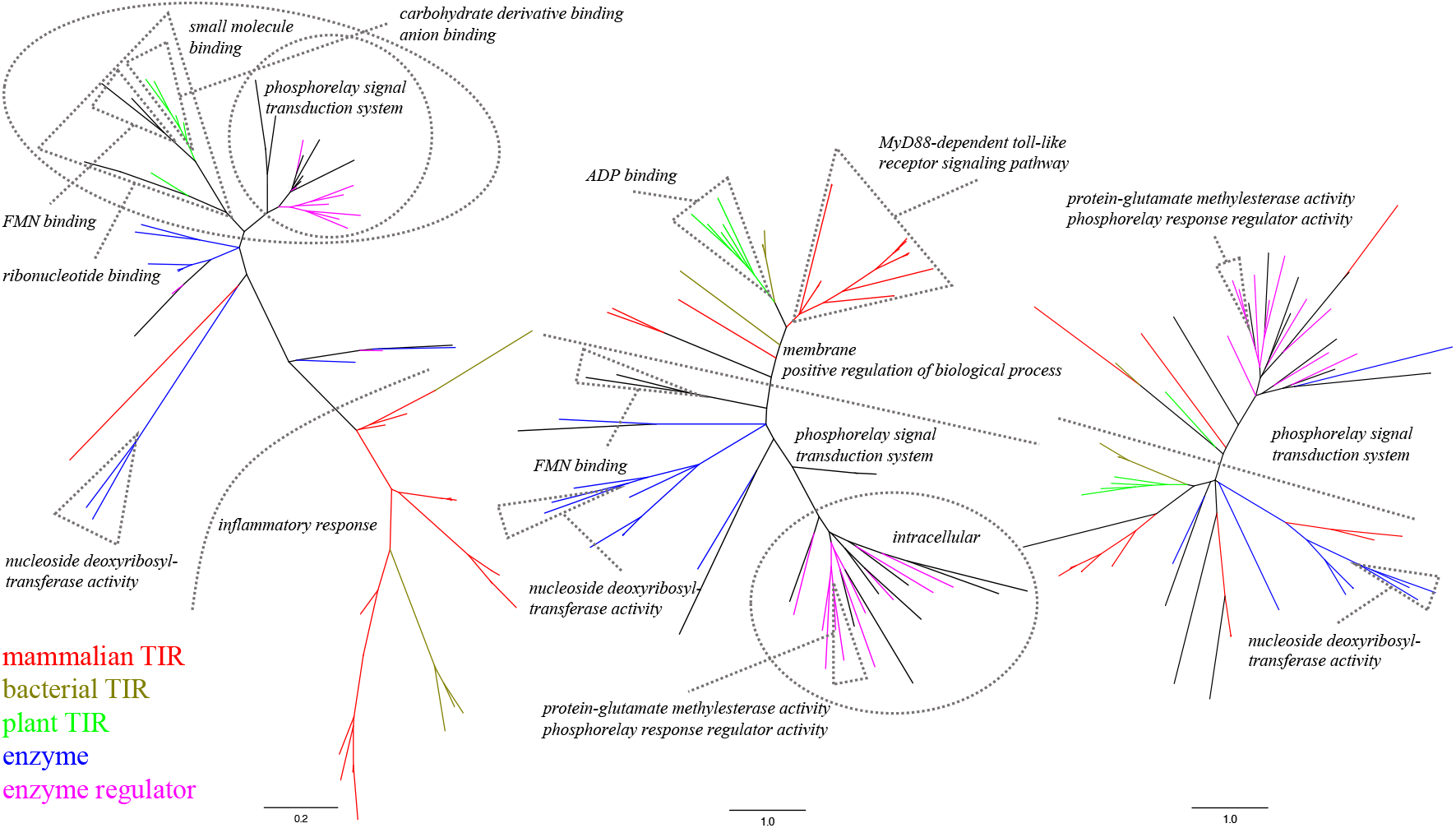
Phylogenetic trees for TIR domains and structurally related proteins based on different evolutionary models and alignments. Left: DSO-sec+jCE; inferred by DSO-sec based on superpositions determined by jCE. Centre: JTT+jCE; inferred by JTT based on the jCE alignment. Right: JTT+MAFFT; inferred by JTT based on an alignment determined by MAFFT. Branch lengths follow the proportions 0.2:1.0:1.0, respectively. Sub-trees with enriched GO terms have been labelled, and groups of proteins are distinguished by colour. We only show the most specific GO term when several are significant. Fully annotated trees are available in SI Figures 2-4.

To qualify how well the trees recovered known biological properties and to quantify their complementarity, we conceived a Gene Ontology (GO) enrichment analysis, which was performed on each tree. Specifically, we determined the statistical over-representation of GO terms for every possible sub-tree, relative to the full tree. The trees created from jCE alignments both highlight evolutionarily meaningful relationships, consistent with our current knowledge of TIR domains and related proteins; these are not apparent in the tree derived from amino acid-based alignment. In JTT+MAFFT, mammalian TIR domains involved in immunity pathways are not well-grouped, nor are proteins with known enzyme activities; only three sub-trees are assigned significant GO terms.

DSO-sec+jCE identifies two important clades: mammalian and bacterial TIR domains versus plant TIR domains, enzymes and enzyme regulators. Based on the same superposition, the secondary structure model infers different evolutionary relationships compared to that based on amino acids. First, in contrast to the amino acid-based tree, the bacterial TIR domains are positioned between mammalian receptor and adaptor TIR domains in the secondary structure-based tree. This is consistent with the finding that several bacterial TIR domain-containing proteins interfere with TLR signalling pathways (Rana et al., 2013). As our analysis showed that bacterial TIR domains have evolved away from both human receptors and adaptors, they may not function as simple mimics of these host proteins. Second, the plant TIR domains are closer to enzyme-related domains than bacterial TIR domains. The structure that is closest to plant TIR domains within the DSO-sec+jCE tree is the flavodoxin-like domain from an oxidoreductase. This domain contains a flavin mononucleotide and is similar to other flavodoxins (Frazão et al., 2000), suggesting that plant TIR domains have enzymatic functions.

The trees in Fig. 2 were each the consensus result of “bootstrapping” 100 trees. To determine the extent with which biological properties were associated with an evolutionary model, each sample tree was evaluated for enriched GO terms; for each model, we ranked each term by the number of times it was deemed significant. A number of terms were equally shared between the models, but a few were predominately associated with either DSO-sec+jCE or JTT+jCE. (JTT+MAFFT is not discussed as it had small counts.) For example, “inflammatory response” was ranked highly (9th) by DSO-sec, and lowly by JTT (44th). Despite being a quite specific term “MyD88-dependent toll-like receptor signaling pathway” was ranked highly by JTT (13th), but lowly by DSO-sec (35th). Both terms are associated with mammalian TLRs but the DSO-sec trees predominately group the bacterial TIR domains with other mammalian TIR domains such that they share a close common ancestor, whereas JTT trees tend to mix in other groups into their common ancestor, which is placed further back. Counts for each tree and each term are found in Additional Data Table S4.

We next tested the utility of the approach on a wider selection of different proteins for which multiple structures were known, and evaluate the differences featured by the evolutionary model of secondary structure when used for phylogenetic analysis, compared to an amino acid-based model. This analysis included the ten largest protein families defined by Pfam with links to the PDB. The average number of structures in each family is 98. For each family, we nominated a “seed” structure; this seed was input to DALI (Holm and Laakso, 2016), which in turn, outputs superpositions of matching structures, that then allowed them to be combined into a multiple sequence alignment.

We performed phylogenetic analyses of these alignments, represented by amino acids and DSSP-defined secondary structure labels, using the JTT and DSO-sec models, respectively. Following the example of TIR-domains, each phylogenetic tree was subjected to a GO enrichment analysis. From this, we note that for nearly all trees, (1) the amino acid model based on the DALI superposition supports a greater number of GO terms, and (2) the secondary structure model finds terms that overlap and complement those of the JTT model. This complementarity is manifested in terms that tend to be relevant to structure, e.g. binding, receptor and enzyme activity (see ABC transporters/PF00005, protein kinase domains/PF00069, reductase/PF13561 and response regulator receiver domains/PF00072 in Additional Data Table S6). In two of ten cases, no GO terms were identified by any method (BPD transporters/PF00528 and Histidine ATPases/PF02518). In one additional case, the model of secondary structure did not identify a single GO term (AMP-binding enzyme/PF00501). These three families have very limited numbers of high-resolution structures (8, 10 and 13 for PF00528, PF02518 and PF07690, respectively). Many GO terms are annotated using sequence similarity. Therefore, sequence-based enrichment studies are to some extent circular. In contrast, our secondary structure based GO enrichment is free of this bias.

To understand structural similarities between evolutionarily distant proteins, exemplified by TIR domains and the ten families from Pfam, we suggest that the set of protein structures is an important source of information, and that they should be looked at as a result of evolution. The next section highlights that, even if we reduce proteins to linear sequences of discrete states, Evolu-sec has the capacity to highlight properties not readily available from modern structures alone.

### 2.3. Reconstructing ancestral secondary structure from modern protein structures

We explored three independent studies to demonstrate how well Evolu-sec (both DSO-sec and JTT-sec) captures evolutionary changes of structural traits. The studies targeted an ancient protein kinase (Wilson et al., 2015), an ancient polar amino acid-binding protein (Clifton and Jackson, 2016), and an ancient DNA-binding domain (Hudson et al., 2016); the proteins represent a range of sequence and structural diversity. Ancestral sequences were inferred from the amino acids of modern proteins, synthesised and their three-dimensional structures were determined. We inferred the secondary structure states of the ancestors using the structures available for a subset of the modern proteins, and evaluated them as described below.

Given the amino acid sequence of an ancestral protein, at least two basic approaches can predict its secondary structure, both of which take as input the amino acid sequence: (1) using secondary structure predictors trained on the wealth of protein structure data, or (2) from the three-dimensional models obtained by comparative modelling techniques referencing databases such as the PDB. Both of these two approaches have been optimised to determine structure from sequence; we would expect that the task would be the same for modern as for ancestral proteins. However, neither approach takes directly and fully into account the evolutionary relationships that must have be known to reconstruct the ancestor in the first place. We wanted to test if Evolu-sec has the capacity to enrich such predictions with evolutionary information.

While our evolutionary model was not expressly designed to predict secondary structure from an amino acid sequence, it can make this prediction for any node in a phylogenetic tree, if secondary structure states are made available at the leaves of such a tree for at least a subset of the leaves. In fact, we do not need the amino acid sequence at all. We used maximum likelihood to find the most probable ancestral secondary structure assignment at the ancestral node, given the corresponding assignments on the leaves of the tree, and the tree itself (Felsenstein, 2003). Ancestral protein structures were available together with sequences and phylogenetic trees; we evaluated the three prediction approaches by comparing their predicted secondary structures with those calculated from experimentally determined high-resolution crystal structures.

In total, our benchmark consisted of seven ancestral protein structures (see Table 2 for details of the data). In each of the three studies, hun­dreds of homologous proteins were used to reconstruct the ancestral protein sequences; we found that only about 10% of the modern proteins used in the studies have experimentally determined structures. We used SWISS-MODEL (Biasini et al., 2014) and I-TASSER (Yang et al., 2014) to perform comparative modelling and we used PredictProtein (Yachdav et al., 2014), JPred (Drozdetskiy et al., 2015) and NetSurfP (Petersen et al., 2009) as representatives of sequence-based predictors.

**Table 2:** Data used for ancestral secondary structure reconstruction.

|  | Wilson et al. | Hudson et al. | Clifton & Jackson |
| --- | --- | --- | --- |
| Modern sequences | 76 | 234 | 340 |
| Mean sequence id. % | 44 | 68 | 30 |
| Modern structures | 10 | 7 | 6 |
| Mean RMSD Å | 1.86 | 0.93 | 1.76 |
| Mean $t$ | 1.66 | 2.31 | 2.94 |
| Ancestor PDB IDs | 4CSV | 5CBX | 4ZV1 |
|  | 4UEU | 5CBY | 4ZV2 |
|  |  | 5CBZ |  |

To calculate the accuracy, we compared predictions made using Evolu-sec on the phylogenetic tree provided in each study (with their evolutionary distances); the 7-class secondary structure was based on the output of comparative modelling (Fig. 3). Because comparative modelling takes each modern protein as the template and produces multiple structures, we present the mean accuracy over all of them. We also compared Evolu-sec based predictions mapped to the 3-class secondary structure with the predictions from PredictProtein, JPred and NetSurfP (Fig. 4; these methods only use 3 out­put classes).

**Figure 3:**
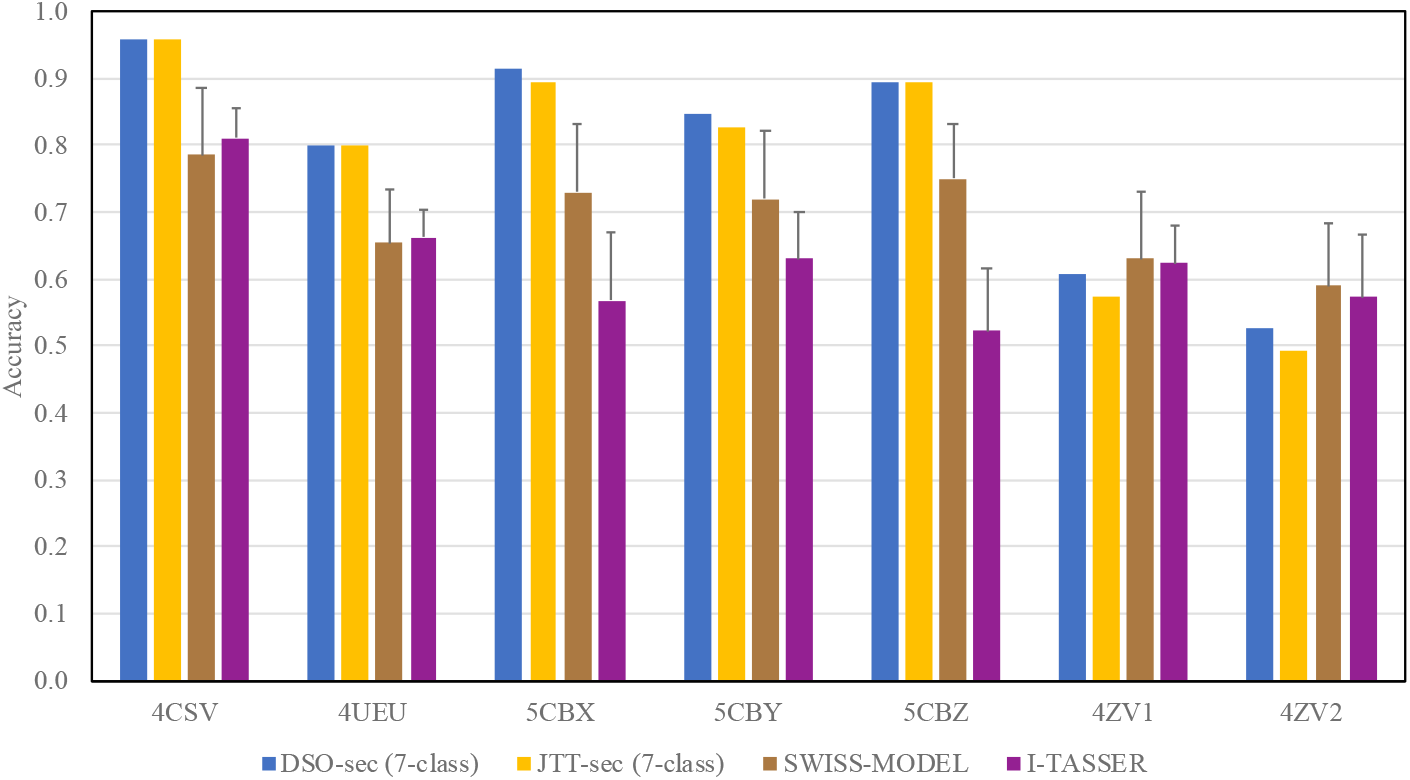
Comparison of 7-class secondary structure prediction accuracy between ancestral secondary structure inferences and comparative modelling (using SWISS-MODEL and I-TASSER). Ancestral secondary structure inferences are based on the secondary structure evolutionary model (DSO-sec and JTT-sec), the original, published amino acid phylogenetic tree, and the available structure data for modern proteins (see text). Homology modelling was performed using each of the modern protein structures and then assigning secondary structure from the models by using DSSP. We averaged the accuracy of comparative modelling-based predictions for ancestral structures; each modern structure is a template for the ancestor.

**Figure 4:**
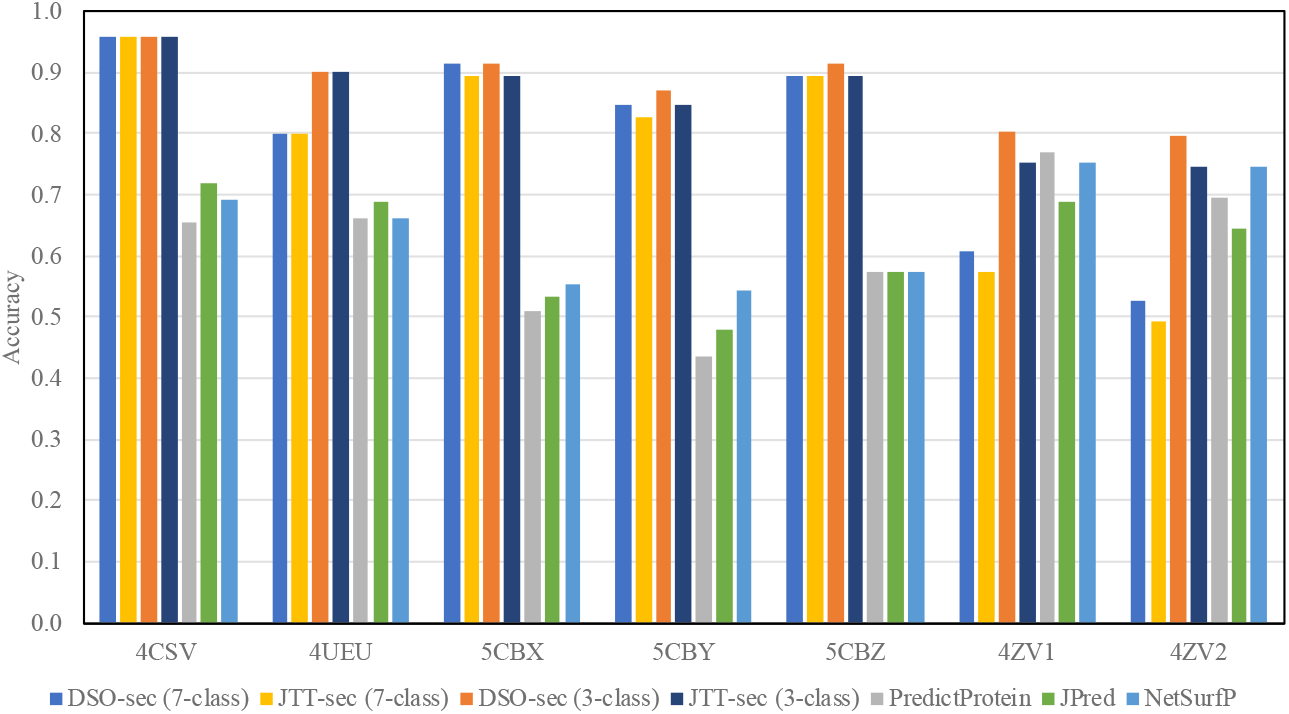
Comparison of 3-class secondary structure prediction accuracy between ancestral secondary structure inferences and sequence-based secondary structure predictions. Predictions were made by state-of-the-art predictors, based on ancestral protein sequences. The 3-class secondary structure annotation corresponds to beta sheet (includes the 7-class B and E), helix (includes the 7-class G, H and I), and other (includes the 7-class S and T).

Evolu-sec based prediction outperformed the accuracy of comparative modelling for 5 of 7 protein structures by 15% or over; it achieved comparable accuracy for the other 2 structures (4ZV1 and 4ZV2, Fig. 3). Across all the proteins, the best accuracy comparative modelling achieved is at 79%, which demonstrates that modern protein structures by themselves are inadequate. Homology modelling did not capture information across multiple, homologous proteins, whereas Evolu-sec explicitly accounted for evolutionary change implied by their placement in the tree.

The accuracy increased when measurements were based on the lower-resolution, 3-class secondary structure definition (Fig. 4). Still, Evolu-sec based prediction outperformed all sequence-based predictors, and the accuracy for 4ZV2 improved from 53% to 80%. The performance on the 3-class definition implies that subtle secondary structure evolutionary changes cannot be distinguished well, e.g. B vs. E.

A few observations are pertinent to understand the capacity of the evolutionary model to capture changes to structure. First, the number of available modern structures did not appear to be a major factor. The worst accuracy was obtained in the dataset with the fewest (6) modern structures; the best average accuracy was obtained in the dataset with 7 modern structures. Second, the single best accuracy was obtained for the ancestral protein structure with modern proteins exhibiting the greatest structural variation (as determined by RMSD); therefore, 3-dimensional dissimilarity did not appear to adversely impact inference.

While sequence-based predictors are known to benefit greatly from capturing the correlative structure between consecutive states, predictions using Evolu-sec are based on observations taken out of their sequence context. We used 19 protein families each with at least 20 members from Pfam to verify that Evolu-sec nevertheless recapitulates a significant tendency to mutate more often at boundaries of, rather than within stretches of secondary structure states (details in SI).

## 3. Discussion

This paper makes available a simple but unique evolutionary model of secondary structure; it offers a platform for pursuing a number of open problems in protein science, which we discuss below. Importantly, we are not arguing that structural traits are the direct subjects of mutation; we do not model changes to the genetic material, but changes to traits that are *accepted* by processes of natural selection. Like Dayhoff’s ground-breaking work, our modest extension is blind to many point mutations in the DNA, including the entire spectrum of “silent” changes (e.g. synonymous codon mutations).

In this paper, we looked at problems that illustrate the potential of integrating protein (secondary) structure into evolutionary analyses. At the outset, we did not expect that pure secondary structure states was a better descriptor for modelling evolutionary change than amino acid composition. Instead, and before combining structural descriptors with traditional sequence states (e.g. nucleic or amino acids), we wanted to highlight if any contribution was evident from secondary structure in its own right.

With Evolu-sec, we were able to infer ancestral secondary structure better than comparative modelling does, provided that protein structures are available for multiple modern proteins. With secondary structure predictors at accuracies above 80% (Drozdetskiy et al., 2015) it is surprising that the outputs of same predictors were inferior in accuracy compared to Evolu-sec reconstructed assignments. The two results demonstrate that Evolu-sec brings together sequence and phylogeny; it captures something about biological structure in addition to what can be recovered from amino acid sequence alone, and in addition to what homology models represent.

Considering that Evolu-sec provides an alternative approach to investigate phylogeny for remote, sequence-diverse, but structurally similar proteins, we explored it for phylogenetic tree inference. Members of the Toll/interleukin-1 receptor (TIR) domain family are involved in innate immunity pathways and are present across plants, mammals, and bacteria; our tests on TIR domains suggests that Evolu-sec detects evolutionary relationships between protein structures that are biologically meaningful but not discerned consistently at the amino acid level.

The standard approach for comparing protein structures revolves around variations of RMSD that determines a geometric fit of structures. The RMSD-based superposition in turn creates an opportunity to create a structural alignment, placing combined information of amino acids so that standard sequence comparison metrics can be used to evaluate transitions. RMSD can be seriously affected by regions outside the common structural core; there is a number of methods based on flexible and elastic structure comparison to make sure that RMSD-based fitting is done only of truly homologous parts (Hasegawa and Holm, 2009). This approach is computationally expensive, and options are sought-after. The present work suggests that Evolu-sec can act as part of an efficient method to enrich a structural similarity “profile” with evolutionary information; we also note that by comparing it with evolutionary distances of an amino acid model it can flag when sequences are not homologous. Evolu-sec places less of a computational burden to perform structural comparison than those based on geometric data; it requires that data are available in the form of linear secondary structure states and that phylogenetic relationships are known. We make available a set of tools to infer phylogenetic relationships including full trees straight from secondary structure assignments.

As flagged in the introduction, we are interested in understanding how we can guide phylogenetic analyses among evolutionarily distant homologues; remote homologues are challenging because they likely display significant diversity at the level of amino acid sequence. We propose that there is a biologically significant complementarity between the evolutionary signal recovered from amino acids and that of secondary structure. We expect that incorporating Evolu-sec as an independent module into ancestral reconstruction pipelines, e.g. by running inference on both types of models in parallel, can help identify sequence targets for protein engineering, e.g. compositionally diverse sites that do not disrupt structural building blocks. We further expect that jointly considering amino acids and secondary structure states will yield an even more accurate instrument; however, to model 140 states (7 secondary structure classes times 20 amino acids), we require more data than what the PDB currently offers.

## 4. Methods and Materials

Methods, data sets and evolutionary models are described in the Supporting Information. In addition, MATLAB scripts to replicate simulations, all data sets and additional figures are available at http://bioinf.scmb.uq.edu.au/evolusec.

## 5. Acknowledgments

The authors thank Dr Yosephine Gumulya for her advice on available ancestral structures. This work was supported by Australian Research Council (ARC) Discovery Project (grant number DP160100865). Bostjan Kobe is a National Health and Medical Research Council (NHMRC) Principal Research Fellow (grant number 1110971).

## Supporting information

Data Table S1: Alignments used for DSO-sec

Data Table S2: Alignments used for JTT-sec

Data Table S3: Evolutionary distances for pairs in Pfam

Data Table S4: Phylogenetic analysis of TIR domains

Data Table S5: Structures in phylogenetic analysis of TIR domains

Data Table S6: Phylogenetic analysis of 10 protein families

