## Supporting Information for "Evolutionary model of protein secondary structure capable of revealing new biological relationships"

### 1. Software and data sets

MATLAB scripts to replicate simulations, all data sets and additional figures are available at `http://bioinf.scmb.uq.edu.au/evolusec`.

### 2. Constructing Evolu-sec by Dayhoff's approach: DSO-sec

We took two steps to create a dataset to model secondary structure changes within homologous proteins.

First, we retrieved all protein crystal structures with a resolution <4.0 Å from the PDB. Secondary structure was assigned by DSSP (Kabsch and Sander, 1983); each residue was assigned one of seven states, i.e., B (beta-bridge), E (beta-strand), G (helix-3), H (alpha-helix), I (helix-5), S (bend), and T (turn), except those residues for which DSSP failed. Peptides or incomplete proteins with fewer than 30 DSSP-assigned secondary structure sites were removed from further consideration. Amino acid sequences for all remaining protein structures were retrieved from UniProt (UniProt Consortium, 2015) and clustered by CD-HIT (Fu et al., 2012) at 85% amino acid sequence identity (as per Dayhoff's original criterion). Clusters with less

*Corresponding author

than 5 proteins were excluded. For each protein accession, when multiple structures were available, we chose the secondary structure from the structure with the best resolution and the greatest number of DSSP-recognised states. The final dataset has 75 clusters, collectively containing 592 proteins. We excluded all structures for reconstructed, ancestral proteins.

Second, we constructed a maximum parsimony tree (Schliep, 2011) based on a multiple sequence alignment generated with MAFFT (Katoh and Standley, 2013) for each cluster. Each alignment consisted of the sequences for which structures are available, and was further enriched with sequences with at least 85% sequence identity from the UniRef90 database; this increased the cluster sizes 20-fold. All trees were subsequently pruned to keep only the tips for which structures are available.

We followed Dayhoff's approach to count evolutionary changes by observing homologous ancestor:descendant pairs of labels (Dayhoff et al., 1978).

Three parameters define the model, namely the accepted point mutation matrix $A$ (shown in Table 1), normalised frequencies ($nFreq$) and relative mutabilities ($rMu$) (shown in Table 2). 1 PAM and 100 PAM are shown in Table 3 and Table 4, respectively. $A_{i,j}$ refers to the number of (undirected) evolutionary changes between $i$ and $j$ that were observed across all sequence positions and all sequence clusters. We excluded sites for which the secondary structure is unassigned.

$MSA^c_{mod}$ is the multiple sequence alignment of modern proteins in cluster $c$. $Tree^c$ is the maximum parsimony tree of $MSA^c_{mod}$. $MSA^c_{anc}$ is the multiple sequence alignment for ancestral proteins (removed from modern protein by one level only), derived from the $Tree^c$ and $MSA^c_{mod}$. $MSSA^c_{mod}$ is the multiple secondary structure alignment, formed from $MSA^c_{mod}$ by replacing amino acid labels with the corresponding secondary structure labels. $MSSA^c_{anc}$ is the multiple secondary structure alignment, formed from ancestral secondary structure labels determined using $Tree^c$ and $MSSA^c_{mod}$. $MSSA^c_{mod}(m,n)$ and $MSSA^c_{anc}(m,n)$ are the two secondary structure labels for protein $m$ in cluster $c$ at sequence position $n$, in its modern and ancestral form, respectively.

We followed (Dayhoff et al., 1978) to calculate the relative mutability $rMu_i$ for a label $i$ as defined in Equation 1. Positions that contain a gap are excluded. The $rMu_i$ is calculated by dividing the total difference exposed to label $i$ with the total occurrence of label $i$. The numerator for $rMu_i$ is to count the total difference for label $i$ in $MSSA^c_{mod}(m,n)$ and $MSSA^c_{anc}(m,n)$ exclusively (Equation 1). The denominator for $rMu_i$ is to count total occurrence for label $i$ in both $MSSA^c_{mod}$ and $MSSA^c_{anc}$ (Equation 1).

$$rMu_i \propto \frac{\sum_c \sum_m \sum_n \begin{cases} 1 & MSSA^c_{mod}(m,n) = i \quad \oplus \\ & MSSA^c_{anc}(m,n) = i \\ 0 & \text{otherwise} \end{cases}}{\sum_c \sum_m \sum_n \begin{cases} 2 & MSSA^c_{mod}(m,n) = i \quad \wedge \\ & MSSA^c_{anc}(m,n) = i \\ 1 & MSSA^c_{mod}(m,n) = i \quad \oplus \\ & MSSA^c_{anc}(m,n) = i \\ 0 & \text{otherwise} \end{cases}} \tag{1}$$

$A^c$ contains the number of observed (un-directional) evolutionary changes between labels $i$ and $j$ when $i \neq j$, for a cluster $c$. We define $A = \sum_c A^c$. $S^c_i$ contains the number of times a label $i$ appears in both the modern and ancestral sequences, for a cluster $c$. We define the frequency of the label $i$ in a cluster $c$ by Equation 2.

$$Freq^c_i = \frac{\sum_k \sum_{l,l \neq k} A^c_{k,l}}{\sum_k \sum_{l,l \neq k} A^c_{k,l} + \sum_k S^c_k} \sum_j A^c_{i,j} \tag{2}$$

Let $nFreq_i = \sum_c Freq^c_i / \sum_i \sum_c Freq^c_i$. $\lambda$ is defined by $nFreq$ and $rMu$ (see Equation 3).

$$\lambda = \frac{0.01}{\sum_i nFreq_i \cdot rMu_i} \tag{3}$$

The non-diagonal elements in a *transition probability matrix* are defined by

$$M_{0.01}(j|i) = \lambda A_{i,j} \frac{rMu_i}{\sum_{k,k \neq i} A_{i,k}} \tag{4}$$

and diagonal elements as

$$M_{0.01}(i|i) = 1 - \sum_{k,k \neq i} M_{0.01}(k|i). \tag{5}$$

Ancestral and descendant states are $i$ and $j$, respectively. $\lambda$ is calibrated to one accepted point mutation per 100 residues ($t = 0.01$), or 1 PAM for short. Dayhoff's original approach lacks precision when extended to longer timeframes, we followed the convention introduced by Kishino et al. (Kishino et al., 1990) and used eigenvalue decomposition to derive the (instantaneous) transition rate matrix $Q$ from $M$ of 7-class secondary structure. Subsequently, we define $M_t$ for any $t$ based on $Q$ (Kosiol and Goldman, 2005).

$$M_t = \exp(tQ) = I + tQ + \frac{(tQ)^2}{2!} + \frac{(tQ)^3}{3!} + ... \quad (6)$$

The $A$ matrix of DSO-sec indicates only 83 evolutionary exchanges between helix (G or H or I) and beta sheet (B or E). The $rMu$ of beta-strand (E) and alpha-helix (H) show they are the most stable secondary structures. This supports the general idea that helix and beta sheet are useful elements for detecting protein homology, because they are unlikely to exchange over short periods. The DSO-sec 1 PAM shows that the Helix-5 (I) secondary structure has the highest probability to mutate to the other states; high relative mutability of Helix-5 suggests an important role in structural evolution (Lee et al., 2000; Fodje and Al-Karadaghi, 2002; Weaver, 2000). The DSO-sec 100 PAM displays a significant deviation in probabilities for all states; the exchangeability between beta sheet and helix increases with a longer distance.

To evaluate the adequacy of the data that underpin DSO-sec, we compared the amino acid evolutionary models DSO and DSO-aa, which are based on Dayhoff's original dataset and our dataset from the PDB, respectively (shown in Table 5). The Pearson correlation for $nFreq$ based on the two datasets is 0.91, and the correlation for $rMu$ is 0.86. The correlation between DSO 1 PAM and that of DSO-aa is 0.99 *with* self-transitions (diagonal elements) and 0.89 *without* self-transitions. The high correlation between the two 1 PAMs suggests that the proteins that we use from the PDB display sequence diversity on par with Dayhoff's original dataset and that data are presented in alignments of adequate quality.

### 3. Constructing Evolu-sec by Jones, Taylor and Thornton's approach: JTT-sec

Jones et al. (Jones et al., 1992) created the JTT model based on an approach that draws more heavily on all protein data available. Accepted point mutations are extracted directly from pairwise alignments; the method only makes reference to modern protein sequences, obviating the need for

| | B | E | G | H | I | S | T |
|---|---|---|---|---|---|---|---|
| B | | 14940 | 140 | 470 | 20 | 3920 | 1220 |
| E | 595 | | 590 | 870 | 0 | 16680 | 2420 |
| G | 0 | 63 | | 22250 | 20 | 11130 | 50260 |
| H | 0 | 20 | 1215 | | 610 | 16680 | 85360 |
| I | 0 | 0 | 0 | 60 | | 30 | 550 |
| S | 158 | 937 | 518 | 663 | 0 | | 80570 |
| T | 33 | 224 | 2575 | 4290 | 40 | 3765 | |

Table 1: The $A$ matrix for secondary structure, as calculated from data based on PDB. As the accepted point mutation matrix ($A$) refers to the undirected evolutionary changes, we present $A$ matrices for DSO-sec and JTT-sec as lower and upper triangles, respectively. As we recorded all evolutionary changes including ambiguous ancestral states as fractional changes for DSO-sec, all numbers were multiplied by 10 and rounded. For example, there are 59.5 changes from E to B, and 93.7 changes from S to E for DSO-sec; there are 1494 changes from E to B, and 1668 changes from S to E for JTT-sec.

| | $nFreq$ | | $rMu$ | |
|---|---|---|---|---|
| | DSO-sec | JTT-sec | DSO-sec | JTT-sec |
| B (Beta-bridge) | 0.0101 | 0.0125 | 460 | 495 |
| E (Beta-strand) | 0.2640 | 0.3032 | 38 | 38 |
| G (Helix-3) | 0.0560 | 0.0459 | 551 | 543 |
| H (Alpha-helix) | 0.4282 | 0.4007 | 100 | 100 |
| I (Helix-5) | 0.0001 | 0.0002 | 7008 | 1526 |
| S (Bend) | 0.1003 | 0.1033 | 391 | 382 |
| T (Turn) | 0.1413 | 0.1342 | 488 | 493 |

Table 2: Normalised frequencies ($nFreq$) and relative mutabilities ($rMu$) for secondary structure. The value for alpha-helix has been arbitrarily set to 100. A larger value in $nFreq$ and $rMu$ means more observations of a structural state and less evolutionarily stable secondary structure, respectively.

| | B | E | G | H | I | S | T |
|---|---|---|---|---|---|---|---|
| B (Beta-bridge) | 9766 | 177 | 0 | 0 | 0 | 47 | 10 |
| E (Beta-strand) | 6 | 9981 | 1 | 0 | 0 | 10 | 2 |
| G (Helix-3) | 0 | 4 | 9720 | 78 | 0 | 33 | 165 |
| H (Alpha-helix) | 0 | 0 | 10 | 9949 | 0 | 5 | 35 |
| I (Helix-5) | 0 | 0 | 0 | 2135 | 6442 | 0 | 1423 |
| S (Bend) | 5 | 31 | 17 | 22 | 0 | 9802 | 124 |
| T (Turn) | 1 | 5 | 58 | 97 | 1 | 85 | 9752 |

Table 3: 1 PAM for secondary structure. The values were multiplied by 10,000. 1 PAM defines evolutionary change over a distance $t = 0.01$ in terms of transition probabilities. 1 PAM is interpreted row-wise.

| | B | E | G | H | I | S | T |
|---|---|---|---|---|---|---|---|
| B (Beta-bridge) | 1092 | 6416 | 192 | 571 | 0 | 1018 | 710 |
| E (Beta-strand) | 227 | 8597 | 94 | 237 | 0 | 518 | 327 |
| G (Helix-3) | 41 | 571 | 1282 | 4161 | 1 | 1432 | 2512 |
| H (Alpha-helix) | 16 | 183 | 528 | 7095 | 1 | 685 | 1492 |
| I (Helix-5) | 30 | 379 | 673 | 5933 | 1 | 1058 | 1926 |
| S (Bend) | 112 | 1616 | 736 | 2770 | 1 | 2407 | 2358 |
| T (Turn) | 54 | 704 | 890 | 4164 | 1 | 1627 | 2559 |

Table 4: 100 PAM for secondary structure. The values were multiplied by 10,000. 100 PAM defines evolutionary change over $t = 1$ in terms of transition probabilities. 100 PAM is interpreted row-wise.

| | $nFreq$ | | $rMu$ | |
|---|---|---|---|---|
| | DSO-aa | DSO | DSO-aa | DSO |
| A | 0.0763 | 0.0870 | 100 | 100 |
| R | 0.0473 | 0.0410 | 78 | 65 |
| N | 0.0482 | 0.0400 | 104 | 134 |
| D | 0.0503 | 0.0470 | 82 | 106 |
| C | 0.0207 | 0.0330 | 33 | 20 |
| Q | 0.0383 | 0.0380 | 76 | 93 |
| E | 0.0574 | 0.0500 | 69 | 102 |
| G | 0.0765 | 0.0890 | 49 | 49 |
| H | 0.0247 | 0.0340 | 89 | 66 |
| I | 0.0523 | 0.0370 | 102 | 96 |
| L | 0.0813 | 0.0850 | 45 | 40 |
| K | 0.0577 | 0.0810 | 80 | 56 |
| M | 0.0214 | 0.0150 | 97 | 94 |
| F | 0.0370 | 0.0400 | 47 | 41 |
| P | 0.0454 | 0.0510 | 36 | 56 |
| S | 0.0720 | 0.0700 | 105 | 120 |
| T | 0.0694 | 0.0580 | 87 | 97 |
| W | 0.0157 | 0.0100 | 26 | 18 |
| Y | 0.0387 | 0.0300 | 48 | 41 |
| V | 0.0693 | 0.0650 | 91 | 74 |

Table 5: Normalized frequencies and relative mutabilities for amino acid models (DSO). Normalized frequencies ($nFreq$) and relative mutabilities ($rMu$) for two amino acid models calculated from Dayhoff's original (DSO) and our dataset (DSO-aa). The value of $rMu$ for alanine has been arbitrarily set at 100.

| | $nFreq$ | | $rMu$ | |
|---|---|---|---|---|
| | JTT-aa | JTT | JTT-aa | JTT |
| A | 0.0847 | 0.0770 | 100 | 100 |
| R | 0.0541 | 0.0510 | 87 | 83 |
| N | 0.0395 | 0.0430 | 123 | 104 |
| D | 0.0571 | 0.0520 | 86 | 86 |
| C | 0.0157 | 0.0200 | 57 | 44 |
| Q | 0.0392 | 0.0410 | 89 | 84 |
| E | 0.0618 | 0.0620 | 89 | 77 |
| G | 0.0798 | 0.0740 | 49 | 50 |
| H | 0.0231 | 0.0230 | 100 | 91 |
| I | 0.0518 | 0.0530 | 122 | 103 |
| L | 0.0866 | 0.0910 | 66 | 54 |
| K | 0.0532 | 0.0590 | 88 | 72 |
| M | 0.0210 | 0.0240 | 128 | 93 |
| F | 0.0379 | 0.0400 | 55 | 51 |
| P | 0.0474 | 0.0510 | 46 | 58 |
| S | 0.0636 | 0.0690 | 135 | 117 |
| T | 0.0629 | 0.0590 | 111 | 107 |
| W | 0.0165 | 0.0140 | 43 | 25 |
| Y | 0.0352 | 0.0320 | 65 | 50 |
| V | 0.0690 | 0.0660 | 110 | 98 |

Table 6: Normalized frequencies and relative mutabilities for amino acid models (JTT). Normalized frequencies ($nFreq$) and relative mutabilities ($rMu$) for two amino acid models calculated from Jones, Taylor and Thornton's original (JTT) and our dataset (JTT-aa). The value of $rMu$ for alanine has been arbitrarily set at 100.

maximum parsimony. The original steps are here extended to secondary structure. We extracted crystal structures from the PDB that with at least 50 amino acids, at resolution 4.0 Å or better, with at least 50% coverage and 40% solved secondary structure (across the whole protein). We used CD-HIT to cluster proteins at 85% and later used MAFFT to perform multiple sequence alignment for all clusters. For each multiple sequence alignment, we extracted pairwise alignment and verified that each pair had a sequence identity of at least 85%. For each protein, we selected one representative structure, giving preference to greater protein coverage and best resolution. Amino acids were then replaced by secondary structure labels determined by the DSSP program. In total, 2,721 pairwise alignments of 2,508 proteins were used to create the secondary structure evolutionary model.

To first benchmark the adequacy of the dataset, we compared the $rMu$ and $nFreq$ values of an *amino acid* model derived from the same 2,508 proteins and found them to be highly correlated with the original JTT at 0.98 and 0.94, with and without self-transitions, respectively. The 1 PAM of our version of JTT and that of the original are correlated at 0.99 and 0.98, respectively.

Using the same 2,508 proteins, we observe 30,873 accepted point mutations in *secondary structure* state. This represents a 20-fold increase of the mutations we used for DSO-sec following Dayhoff's approach (see Table 1). Again, $nFreq$ and $rMu$ of DSO-sec and JTT-sec are highly correlated at 0.99 and 0.94, respectively (shown in Table 6). The 1 PAM of JTT-sec and 1 PAM of DSO-sec are correlated at 0.99 with self-transitions, and 0.90 without self-transitions. We note that the dynamic range of JTT-sec (as per the Kullback-Leibler divergence metric) follows the tendency of DSO-sec with greater sensitivity at shorter time ranges. The IRM for JTT-sec is presented in Table 7.

### 4. Evolutionary distances based on secondary structure are shorter than those based on amino acids

To qualify in relative terms what Evolu-sec does, we investigated evolutionary changes at both the structure and amino acid levels; we probed transition probabilities at different points in time, as implied by Evolu-sec (both DSO-sec and JTT-sec) and the amino acid models (DSO and JTT).

The evolutionary distance ($t$) determines the transition probabilities that apply between any pair of states; at time $t = 0$, the transition probabilities

| | B | E | G | H | I | S | T |
|---|---|---|---|---|---|---|---|
| B (Beta-bridge) | -26641 | 19224 | 173 | 590 | 27 | 5075 | 1552 |
| E (Beta-strand) | 851 | -2005 | 32 | 47 | 0 | 944 | 131 |
| G (Helix-3) | 46 | 196 | -29298 | 7634 | 7 | 3807 | 17607 |
| H (Alpha-helix) | 20 | 35 | 940 | -5344 | 27 | 691 | 3632 |
| I (Helix-5) | 1387 | 0 | 1253 | 41771 | -84436 | 1871 | 38153 |
| S (Bend) | 629 | 2643 | 1752 | 2582 | 4 | -20554 | 12943 |
| T (Turn) | 145 | 276 | 6119 | 10254 | 69 | 9772 | -26635 |

Table 7: Instantaneous rate matrix for secondary structure based on Jones, Taylor and Thornton's approach (JTT-sec). The values were multiplied by 10,000. The instantaneous rate matrix is interpreted row-wise.

are given by the Kronecker delta, and as time approaches infinity, transition probabilities converge to a stationary distribution. To understand the differences between Evolu-sec and amino acid models in terms of transition probabilities at different distances from a reference time $t = 0$, we used the Kullback-Leibler divergence (Kullback and Leibler, 1951); we quantified the rate with which transition probabilities lose their dynamic range and become saturated, over time. We framed this rate as the *average divergence* (for each possible ancestral state $i$) over an interval in ancient time, defined by $t$ and $t - d$ (Equation 7).

$$E_t = \frac{1}{n} \sum_i \sum_j M_t(j|i) \log(\frac{M_t(j|i)}{M_{t-d}(j|i)}) \tag{7}$$

$t$ and $d$ define a time interval away from the reference time, $t > d > 0$; the parameter $i$ is the ancestral state and $j$ is the descendant state, and $n$ is the number of states in the model, i.e. $n = 7$ for 7-class secondary structure and $n = 20$ for the amino acid model; $M_t$ is the transition probability matrix at time $t$.

We calculated the average divergence at offsets from the reference time $0.1 \leq t \leq 1.0$; we used a constant interval $d = 0.09$ over which we determined the divergence (see Fig. 1). For all models, the dynamic range (as quantified via Equation 7) reduced monotonically as we moved back in time, until convergence when the present-time state had no discernable impact on the ancient-time state. This was represented by a negligible divergence between the two ancient time points.

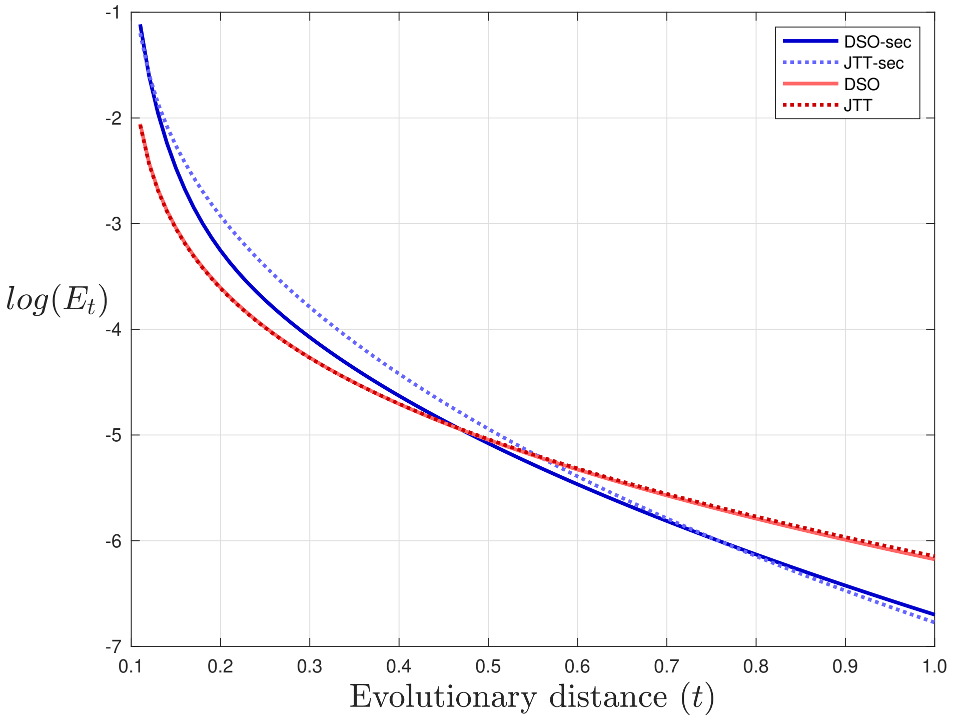


Figure 1: Kullback-Leibler divergence between transition probabilities over a constant distance $d = 0.09$ at different offsets $(t)$.

It is important to note that the divergence metric cannot be used to compare the models in an absolute sense, because the numbers of ancestral character states are different (7 and 20, respectively). However, once they are looked at over the same time axis, a few observations can be made. Relative to ancient time, closer to present time, Evolu-sec displayed a dynamic range of transition probabilities that is greater than that of amino acid models (DSO or JTT); farther offset from the present time, both models converged towards their stationary distributions, which are not sensitive to the state at the reference time. However, Evolu-sec converged more quickly, potentially due to back substitutions inherent from a smaller state space.

### 5. Inferring evolutionary distance and phylogenetic tree

Following Felsenstein's approach (Felsenstein, 1996), the likelihood $\mathcal{L}$ is defined for an evolutionary distance $t$, based on the transition probability matrix $M_t$ as in Equation 8.

$$\mathcal{L} = \prod_i \prod_j (\pi_i M_t(j|i))^{n_{i,j}} \tag{8}$$

The normalised (equilibrium) frequency for the ancestor $i$ is $\pi_i$. $M_t(j|i)$ is the probability of $j$ given ancestor state $i$. $n_{i,j}$ is the observed changes from the state $i$ to state $j$ between two aligned protein sequences. The evolutionary distance $t$ is estimated by maximising the likelihood $\mathcal{L}$. Our implementation of the Equation 8 is to estimate evolutionary distances for secondary structure states. The same implementation in PHYLIP package (Felsenstein, 2005) was used to estimate evolutionary distances for amino acids. The unit of the evolutionary distance $t$ is scaled as the unit of expected number of substitutions per site.

We extracted alignments from the A-seed dataset from the Pfam database version 30.0 (Finn et al., 2016). We excluded sequences with non-standard amino acids. Pfam defines an eighth label C for irregular secondary structure, which we treated as unknown. We created alternative datasets that conform to different percentage cut-offs for assigned secondary structure, specifically at 50%, 60%, 70%, and 80%. In all, we estimated 26,480 pairwise distances within 1,591 domains. The Pearson correlations between estimated distances for sequences that display 50%, 60%, 70%, and 80% of residues with an assigned secondary structure state, are 0.48 (26,291 pairs), 0.51 (25,552 pairs), 0.54 (22,157 pairs), and 0.56 (13,227 pairs), respectively.

### 6. Constructing a secondary structure phylogenetic tree for TIR domains

We developed a phylogenetic tree inference method based on Evolu-sec and maximum likelihood. Our tree inference method broadly follows the approach of PHYML (Guindon and Gascuel, 2003); it uses UPGMA and the Adachi's least square method (Adachi and Hasegawa, 1996) to assign an initial tree; it uses nearest neighbourhood inter-exchange and Brent's algorithms (Brent, 1973) to search the best tree by maximising the tree likelihood.

As the TIR domain from human Toll-like receptor 1 (PDB code 1FYV) is the first crystal structure of any TIR domain, we used Dali server (Holm and Laakso, 2016) to search any target structure data similar to this structure within the PDB. We removed redundant chains within each structure and kept crystal structures with a resolution less than 4.0 Å. As per Dali's recommendation, we used structures with a Z-score less than 2 and a superposition RMSD less than 4 Å. Based on Dali's initial superposition, we only used structured data superimposed onto 1FYV with $> 70\%$ of residues and composed of 125-200 residues. This is to avoid including small fragments and to avoid large insertions within the alignment. In the end, 64 structure data were used for analysis. The multiple structural alignment was performed by jCE (Java combinatorial extension) with default parameters, scoring both C$\alpha$ distances and angles between sidechains (Prlic et al., 2010). A structure-assisted MSA was generated by following the structural alignment procedure described above, after which the amino acid MSA was extracted from the individual protein structures. We used PHYML (JTT model, four categories and maximum likelihood estimated gamma shape parameter) to infer phylogenetic trees from the amino acid MSA derived by superposition and MAFFT. We performed 100 replicates of regular bootstrapping by SEQBOOT and CONSENSE program (part of PHYLIP).

To test significant GO annotations within the clades of a tree, we explored all possible subtrees. As three phylogenetics trees are unrooted, we defined tips of a subtree as the positive set and the remaining tips as the negative set. We used both direct and indirect GO annotations for two sets and used Fisher's exact test to detect significant annotations with $E$-value less than 0.05 (corrected for multiple tests). Three phylogenetic trees, DSO-sec+jCE, JTT+jCE, and JTT+MAFFT are shown in Fig. 2, Fig. 3, and Fig. 4,respectively.

### 7. Phylogenetic analysis of ten large protein families

We extracted ten large protein domain families and associated PDB IDs from Pfam. We started with Pfam's 20 largest families, excluded repeated domains and selected the ten with the longest protein sequences. The average sequence identity within each family is 21%.

We nominated the first published high-resolution structure as the seed, for each domain family. With the following exceptions, we used the same strategy as for the TIR domain tree to construct the phylogenetic tree for

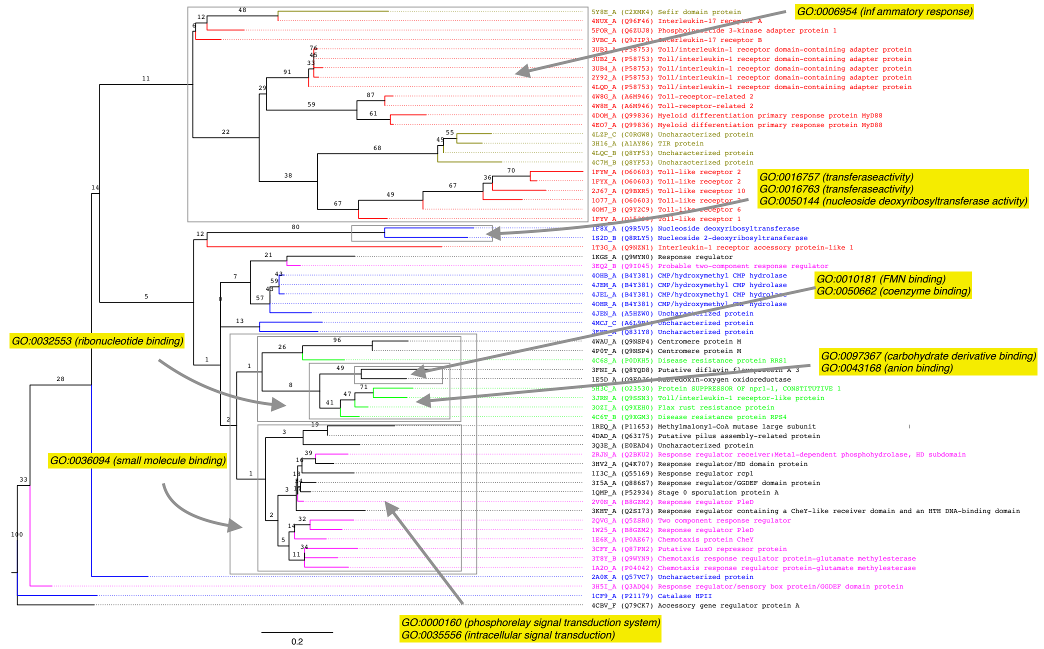


Figure 2: Phylogenetic trees for TIR domains inferred by DSO-sec based on superpositions determined by jCE with GO annotations

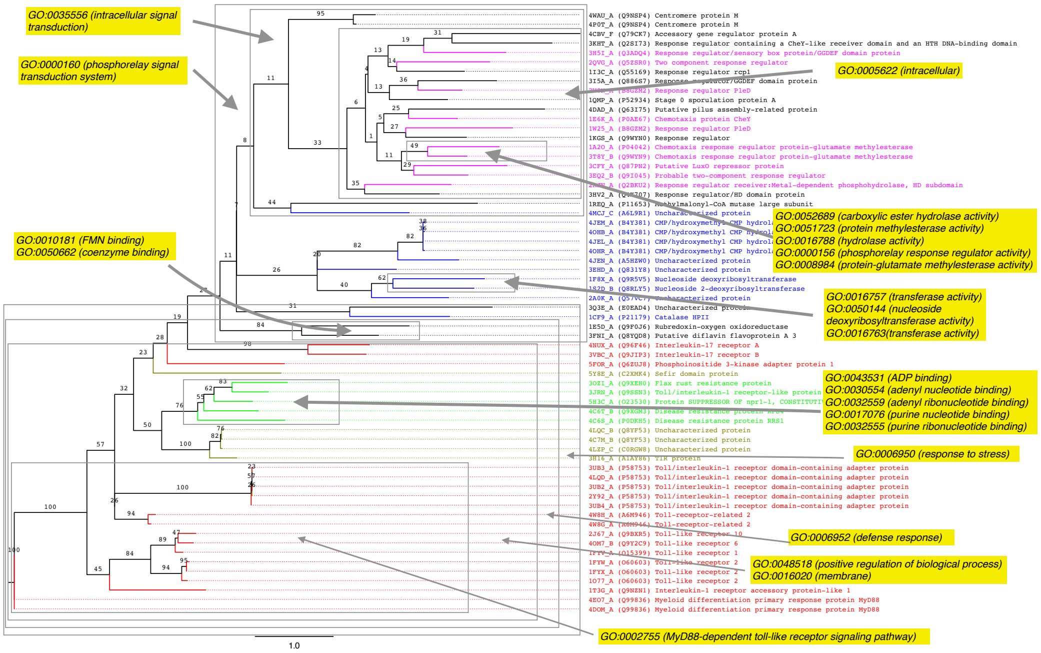


Figure 3: Phylogenetic trees for TIR domains inferred by JTT based on superpositions determined by jCE with GO annotations

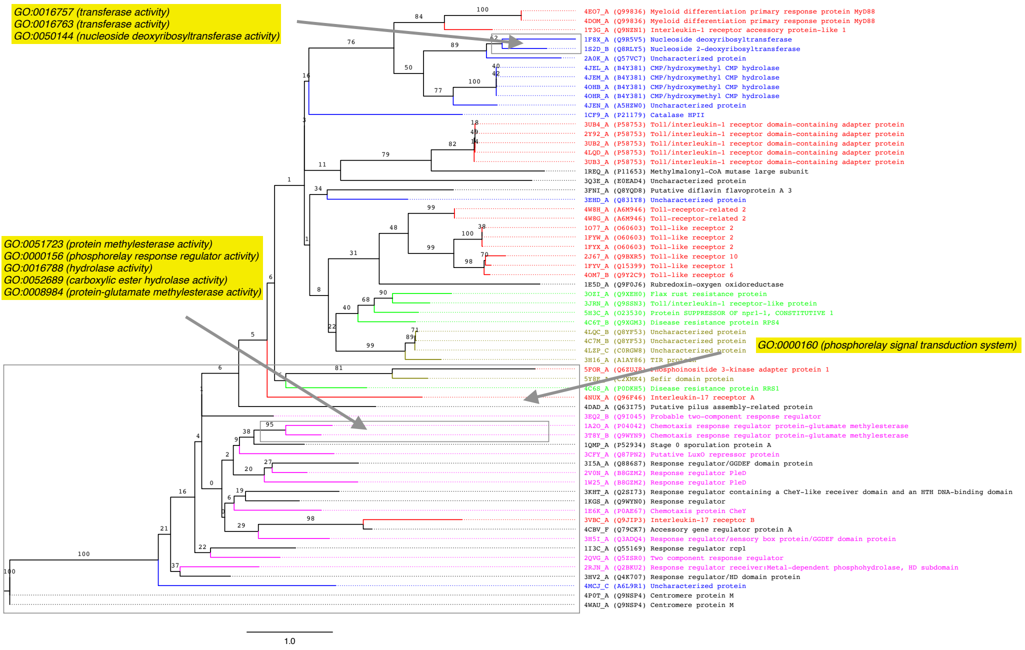


Figure 4: Phylogenetic trees for TIR domains inferred by JTT based on MAFFT with GO annotations

each Pfam domain. First, we searched Dali's redundancy reduced data set with less than 50% sequence identity. Second, the structural alignments were taken from Dali, and not jCE. Third, we did not exclude proteins based on their lengths. While Dali provides 3-class secondary structure labels, we converted them back to the 7-class DSSP labels, by reference to the original PDB records.

### 8. Inferring ancestral secondary structure

Wilson et al. (Wilson et al., 2015), Hudson et al. (Hudson et al., 2016), and Clifton and Jackson (Clifton and Jackson, 2016) inferred, synthesised ancestral proteins and solved one or more of their structures. We retrieved the original phylogenetic tree and the amino acid sequence alignment for each of the three studies. Each tree topology was pruned to only contain tips for which structure data were available. For Wilson et al., the pruned tree contains 10 modern protein structures: 1QCF, 2H8H, 3A4O, 3BYS, 3DTC, 3GVU, 3SXS, 4HCT, 4XEY and 4YHF. The mean pairwise structure RMSD is 1.86 Å and the mean pairwise evolutionary distance is 1.66. For Hudson et al., the pruned tree contains 7 modern protein structures: 1HCQ, 1R4I, 2C7A, 3G9M, 4CN5, 4TNT and 5EMQ. The mean pairwise structure RMSD is 0.93 Å and the mean pairwise evolutionary distance based on an amino acid evolutionary model is 2.31. For Clifton and Jackson, the pruned tree contains 6 modern protein structures: 1HSL, 1WDN, 4I62, 4KQP, 4ZEF and 5EYF. The mean pairwise structure RMSD is 1.76 Å and the mean pairwise amino acid evolutionary distance is 2.94. The RMSD was calculated by jCE with superposition algorithm scoring both C$\alpha$ distances and angles between sidechains.

Multiple secondary structure alignments were created by substituting amino acids for secondary structure states. We determined the ancestral secondary structure with the greatest maximum joint probability from the pruned trees and the alignments, using the Evolu-sec models. Columns with gaps were ignored and normalised (equilibrium) frequencies were assigned to sites with unknown secondary structure state.

We evaluated prediction accuracy by comparing inferred ancestral secondary structure to that determined by DSSP for the actual ancestral structures deposited in the PDB. The 7-class accuracy is defined as the number of correctly predicted sites, divided by the number of sites for which a prediction was made, i.e., the columns of the alignment that have more than one inconsistent variation or unknown secondary structure state. Using the Wilson et al. case as an example, there are 319 columns in the original alignment. We reconstructed only 237 columns by removing 82 columns with one or more gaps from 319 columns in the alignment; we also removed 117 columns with no variation, i.e. identical labels. After this filtering, only 120 columns are used to evaluate the accuracy of reconstruction. Once we consider the ancestral structure, we note that only 75 out of the 120 columns have been assigned secondary structures by running DSSP on 4CSV. A correct prediction is made for 72, meaning the accuracy is 0.96 (72/75).

SWISS-MODEL (Biasini et al., 2014) and I-TASSER (Yang et al., 2014) were used to model the 3D structure by assigning each modern protein structure in the tree as a template; and we assigned secondary structure by DSSP and evaluated 7-class accuracy for each predicted structure. PredictProtein (Yachdav et al., 2014), JPred (Drozdetskiy et al., 2015), and NetSurfP (Petersen et al., 2009) were used to conduct sequence-based predictions; accuracies were recalculated by 3-class secondary structure as helix (G or H or I), beta sheet (B or E), and other (S or T).

### 9. The model sustains correlation between structural states in sequence

Protein conformation depends on sequence context (Anfinsen et al., 1961). It was noticed early on that neighbouring secondary structure states are correlated (Chou and Fasman, 1974). It is also well-established that secondary structure predictors capturing dependencies over a window of positions achieve greater accuracy (Rost and Sander, 1993). Evolu-sec is based only on observations taken out of their context, so it is unclear if it can promote correlation between consecutive residues when used for analysing full-length sequences.

As a straw man test, we expected the level of conservation over full-length sequences to be greater at positions embedded within the same secondary structure state, than those at boundaries of structural domains. We defined a boundary position to be one which is assigned a secondary structure state which is *not* identical to *both* of its neighbours. Any other position is non-boundary, i.e. within a stretch of the same state.

A model of evolution should ideally reproduce sequences with the same correlated structure, so we asked two questions.

1. What is the baseline variability between boundary and non-boundary sequence contexts reflected between modern and actual, reconstructed ancestral protein structures?
2. Is the baseline variability maintained by Evolu-sec, when used to predict ancestral secondary structure?

To answer the two questions, we again used the seven ancestral structures from the previous section. In addition, to further quantify if Evolu-sec maintained correlated secondary structure assignments, we analysed nineteen Evolu-sec inferred trees of Pfam domains. We first grouped positions as either boundary or non-boundary. Then we determined the fraction of cases for each group that represented a substitution of secondary structure state relative its ancestor. This was either done on basis of an available ancestral structure–for question 1, or on basis of an Evolu-sec predicted ancestral secondary structure–for question 2.

Boundary residues are sequence sites that are not surrounded by the same secondary structure state. Ancestral protein structures and their phylogenetic trees from three published works were used to study evolutionary changes at boundaries (Hudson et al., 2016; Wilson et al., 2015; Clifton and Jackson, 2016). Moreover, we used 19 Pfam domains that have multiple secondary structure alignment of more than 20 proteins in the A-seed file. For each domain, we used Evolu-sec to infer a phylogenetic tree; we then inferred ancestral secondary structure, and determined the frequency of change (boundary vs. non-boundary) for each protein. For all our cases, structures were always available for modern proteins; this was not the case for ancestors. To avoid circularity of predicting the ancestral state, we based boundaries on the state within modern proteins.

In regard to the first question, the baseline rate of variation was $4.72 \pm 1.67$ for the seven ancestral structures; this was based on frequencies of mutation at boundaries and at non-boundaries, averaging $0.15 \pm 0.03$ and $0.04 \pm 0.01$, respectively. In other words, the secondary structure was almost five times more likely to be substituted at the boundary of local conformations, compared to when embedded.

In regard to the second question, based on Evolu-sec predictions from on the descendants of the seven ancestral structures, the rate was at a comparable $6.19 \pm 0.76$. The frequencies of mutation at boundaries and non-boundaries averaged $0.20 \pm 0.04$ and $0.03 \pm 0.01$, respectively. The rate was $6.15 \pm 2.98$ over the nineteen Evolu-sec inferred Pfam trees; average mutation frequencies were $0.06 \pm 0.03$ and $0.01 \pm 0.01$ for boundaries and non-boundaries, respectively. Based on Evolu-sec, over full-length sequences, the rate of substitution was about six times more likely at boundaries. Taken together, secondary structure has a clear tendency to mutate at boundaries of secondary structure states, a trend which is reproduced by the evolutionary model. All the results above were based on DSO-sec, but JTT-sec displayed the same trend.

Yachdav, G., Kloppmann, E., Kajan, L., Hecht, M., Goldberg, T., Hamp, T., Hönigschmid, P., Schafferhans, A., Roos, M., Bernhofer, M., Richter, L., Ashkenazy, H., Punta, M., Schlessinger, A., Bromberg, Y., Schneider,

R., Vriend, G., Sander, C., Ben-Tal, N., Rost, B., 2014. Predictprotein–an open resource for online prediction of protein structural and functional features. Nucleic Acids Res 42, W337–W343.

Yang, J., Yan, R., Roy, A., Xu, D., Poisson, J., Zhang, Y., 2014. The I-TASSER Suite: protein structure and function prediction. Nat Methods 12, 7–8.
